# Astrocytes express DMT1 and transferrin receptors, which transport iron thus activating Ca^2+^ signalling: possible role in neuroprotection against iron overload?

**DOI:** 10.1101/2020.07.06.190652

**Authors:** Maosheng Xia, Wenzheng Guan, Ming Ji, Shuai Li, Zexiong Li, Beina Chen, Manman Zhang, Shanshan Liang, Binjie Chen, Wenliang Gong, Chengyi Dong, Gehua Wen, Xiaoni Zhan, Dianjun Zhang, Xinyu Li, Alexei Verkhratsky, Baoman Li

## Abstract

Iron is the fundamental element for numerous physiological functions. Reduced ferrous (Fe^2+^) and oxidized ferric (Fe^3+^) are the two ionized iron states in the living organisms. In the cell membrane, divalent metal ion transporter 1 (DMT1) is responsible for cellular uptake of Fe^2+^, whereas transferrin receptors (TFR) carry transferrin (TF)-bound Fe^3+^. In this study we performed, for the first time, detailed analysis of the action of Fe ions on cytoplasmic free calcium ion concentration ([Ca^2+^]_i_) in astrocytes. Using qPCR and immunocytochemistry we identified DMT1 and TFR in astrocytes in primary cultures, in acutely isolated astrocytes and in brain tissue preparations; *in situ* both DMT1 and TFR are concentrated in astroglial perivascular endfeet. Administration of Fe^2+^ or Fe^3+^ in low μM concentrations evoked Ca^2+^ signals in astrocytes *in vitro* and in *vivo*. Iron ions triggered increase in [Ca^2+^]_i_ by acting through two distinct molecular cascades. Uptake of Fe^2+^ by DMT1 inhibited astroglial Na^+^-K^+^-ATPase (NKA), which led to an elevation in cytoplasmic Na^+^ concentration (as measured by SBFI probe), thus reversing Na^+^/Ca^2+^ exchanger (NCX) thereby generating Ca^2+^ influx. Uptake of Fe^3+^ by TF-TFR stimulated phospholipase C to produce inositol 1,4,5-trisphosphate (InsP_3_), thus trigering InsP_3_ receptor-mediated Ca^2+^ release from the endoplasmic reticulum. Iron-induced Ca^2+^ signals promote astroglial release of arachidonic acid and prostaglandin E2 cytokines by activating cytosolic phospholipase A2 (cPLA2) and NF-κB signalling cascade. In summary, these findings reveal new mechanisms of iron-induced astrocytic signalling operational in conditions of iron overload, in response to which astrocytes actively accumulate excessive iron and activate neuroprotective pathways.

## INTRODUCTION

Iron contributes to numerous cellular and biochemical processes and acts as a co-factor in various molecular cascades in the nervous tissue including the synthesis and metabolism of several brain-specific enzymes and neurotransmitters ^1, 2^. In biological systems iron is present in either reduced ferrous (Fe^2+^) or oxidized ferric (Fe^3+^) state. The brain has the second (after liver) highest quantity of iron in the human body with total non-heme iron in the brain reaching about 60 mg ^3^. The non-heme iron concentration in the serum ranges between 9-30 μM, whereas the iron concentration in cerebrospinal fluid (CSF) is much smaller being around 0.3-0.75 μM ^4, 5^. Transport of iron across the blood-brain barrier (BBB) is mediated either by means of transferrin receptor (TFR)-mediated internalisation of Fe^3+^-bound to transferrin (holo-TF), or, for non-TF-bound iron, by vesicular and non-vesicular pathways ^6^. Membrane transport of Fe^2+^ is also mediated by divalent metal ion transporter 1 (DMT1/SLC11A2) which underlies Fe^2+^ uptake through the plasma membrane or from endosomes ^6^. Under physiological conditions, the intracellular cytosolic ionized iron levels fluctuate around 0.5-1.5 μM ^7^.

In the brain, up to three-fourths of total iron is accumulated within neuroglial cells ^8^. Astrocytes in particular are fundamental elements of ionostatic control over CNS environment ^9^. Ionized Fe^2+^ enters astrocytes through DMT1/SLC11A2 transporters which are particularly concentrated in endfeet of cerebral and hippocampal astrocytes ^10^. The evidence for the expression of TFR in astroglial cells remains controversial ^6, 11, 12^, while iron overload may influence the expression or distribution of TFR in astrocytic compartments ^12, 13^. Cellular uptake of Fe^3+^ requires internalization of TF-TFR complex ^14^. An adaptor protein Disabled-2 (Dab2) plays an essential role in cell signalling, migration and development ^15^. In mice the Dab2 has two isoforms of 96 and 67 kDa (p96 and p67 ^15^). In human K562 cells, Dab2 regulates internalization of TFR and uptake of TF ^16^. Dab2 is also widely distributed in immune cells and in neuroglia ^15^, although the functional link between Dab-2 and TFR in astrocytes has not been demonstrated.

Astrocytes possess a special form of intracellular ionic excitability, mediated by temporal and spatial fluctuations in the intracellular ion concentration ^17, 18^. Astroglial Ca^2+^ signalling is mediated by Ca^2+^ release from the endoplasmic reticulum (ER) following activation of inositol-1,4,5-trisphosphate receptor (InsP_3_R), or intracellular Ca^2+^-gated Ca^2+^ channels known as ryanodine receptors (RyR). Astroglial Ca^2+^ sugnals may also be generated by plasmalemmal Ca^2+^ entry through Ca^2+^-permeable channels or by sodium-calcium exchanger (NCX) operating in the reverse mode ^17, 19^. Astroglial Na^+^ signalling is shaped by plasmalemmal Na^+^ entry through cationic channels and numerous Na^+^-dependent transporters, of which the major role belongs to Na^+^-dependent glutamate transporters ^20–22^; as well as by Na^+^ extrusion through the sodium-potassium pump (NKA). Both Na^+^ and Ca^2+^ signalling systems are closely coordinated, with NKA and NCX accomplishing this coordination at the molecular level ^23^. Astrocytes specifically express α2-subunit containing NKA which is fundamental for astroglial K^+^ buffering ^24^. Astrocytes express all three isoforms of NCX - NCX1/SLC8A1, NCX2/ SLC8A2 and NCX3/SLC8A3, with some evidence indicating higher expression of NCX1 ^25^. The NKA, the NCX and glutamate transporters are known to be preferentially concentrated in the perisynaptic astroglial membranes indicating intimate relationship between these ion-transporting molecules ^26, 27^. The NCX are also known to localise at caveolae with caveolin-3 (Cav-3), the latter isoform being predominantly expressed in astrocytes ^28^.

The excess of iron may cause abnormal [Ca^2+^]_i_ signalling and the dysregulation of the downstream kinase cascades. Iron can increase the protein expression of P65 subunit of nuclear factor κB (NF-κB) ^29^, which was claimed to contribute to neuroinflammation and neuroprotection through activation of cytosolic phospholipase A2 (cPLA2) in astrocytes ^30^. The cPLA2 is a Ca^2+^-dependent phospholipase, which can hydrolyze the fatty acid at sn-2 position of glycerophospholipids to produce arachidonic acid (AA), which is an astroglia-specific process ^31^. In our previous studies we demonstrated that the Ca^2+^-stimulated phosphorylation of cPLA2 contributes to the fast secretion of AA and prostaglandin E2 (PGE2) from spinal cord astrocytes ^32, 33^.

In the present paper we performed an in depth analysis of the action of ferrous and ferric (Fe^2+^ and Fe^3+^) on astroglial Ca^2+^ and Na^+^ dynamics. We found that Fe^2+^ (via DMT1) and Fe^3+^-TF (via TFR) evoke [Ca^2+^]_i_ transients in astrocytes in culture and *in vivo*. Effects of Fe^2+^ on [Ca^2+^]_i_ were mediated mainly by the reversed NCX, whereas Fe^3+^ triggered Ca^2+^ release from the endoplasmic reticulum by stimulations of InsP_3_R. Finally, we quantified iron-induced downstream activity of cPLA2 and NF-κB as well as cPLA2-regulted up-regulation of arachidonic acid (AA) and prostaglandin E2 (PGE2).

## RESULTS

### Fe^2+^/Fe^3+^ trigger [Ca^2+^]_i_ increase in cortical astrocytes *in vitro* and *in vivo*

We analysed Ca^2+^ dynamics in astrocytes in primary cultures and *in vivo* in the GFAP-eGFP transgenic mice for cell identification (Fig. 1). In the primary cultured astrocytes, administration of both FeSO_4_ (Fe^2+^) or ferric ammonium citrate-TF (Fe^3+^) increased [Ca^2+^]_i_ in concentration-dependent manner, albeit with different kinetics. In the presence of Fe^2+^ an increase in [Ca^2+^]_i_ demonstrated prominent plateau, whereas Fe^3+^ triggered transient relatively rapidly decaying [Ca^2+^]_i_ rise (Fig 1A). Both ions induced [Ca^2+^]_i_ responses in a concentration dependent manner with apparent EC_50_ of 0.635 μM for Fe^2+^ and 0.711 μM for Fe^3+^ (Fig. 1C).

**Figure 1.**
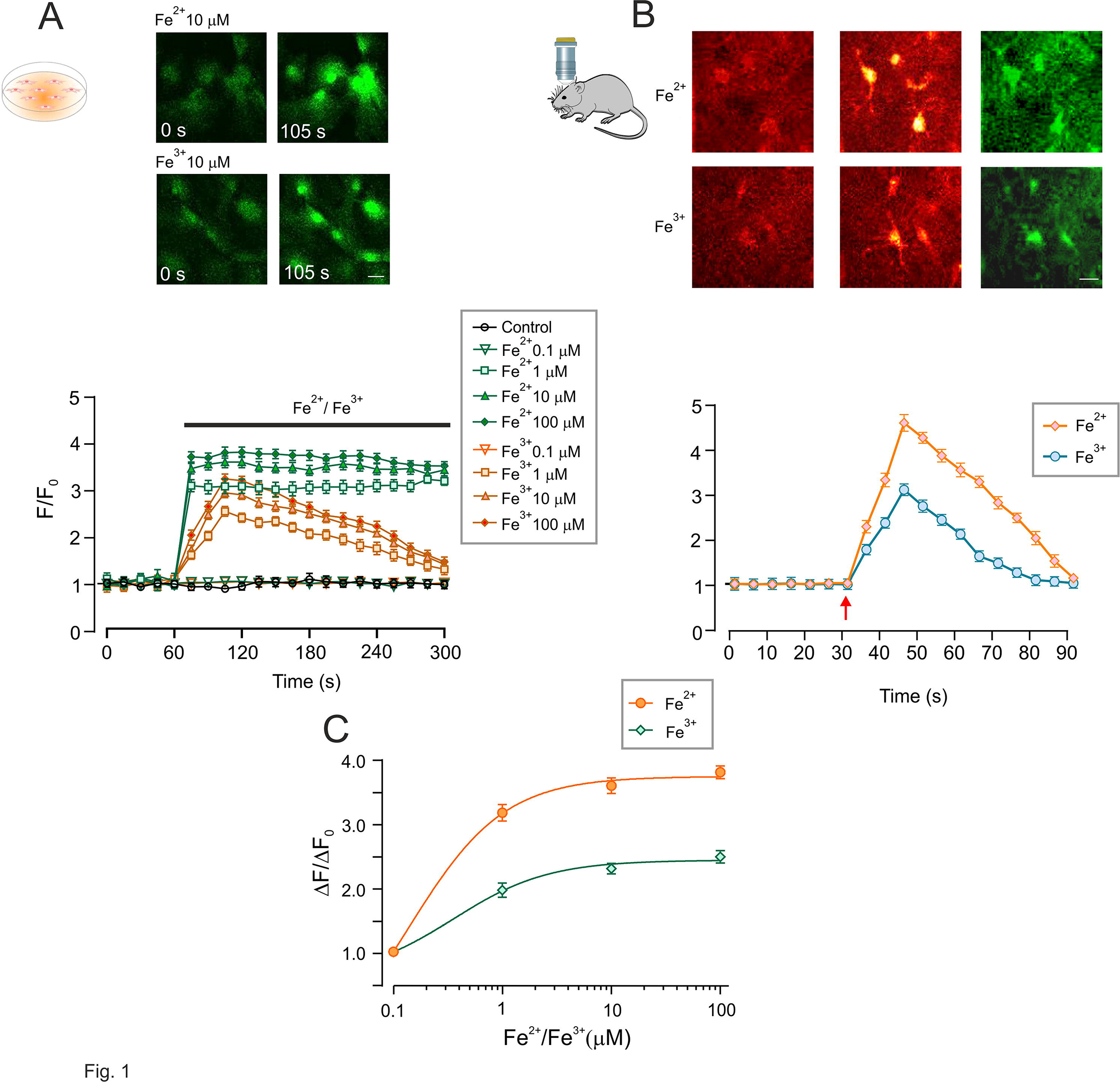
Iron ions, Fe^2+^ and Fe^3+^, evoke astrocytic intracellular Ca^2+^ signals *in vitro* and *in vivo*. (A) Images (top panel) and intracellular Ca^2+^ ([Ca^2+^]_i_) recordings from primary cultured astrocytes in response to different concentrations of Fe^2+^ or Fe^3+^. Every data point represents mean ± SD, n = 10, p < 0.05, statistically significant difference from the value of baseline in the same group. Representative Fluo-4 images show astrocytes treated by 10 μM Fe^2+^ or Fe^3+^ at 0 second and 105 second of the recordings. Scale bar, 20 μm. (B) Images (top panel) and [Ca^2+^]_i_ recordings from cortical astrocytes in GFAP-eGFP transgenic mice using transcranial confocal microscopy. Every data point represents mean ± SD, n = 10. Representative Rhod-2 images (red) show astrocytes treated with 100 μM Fe^2+^ or Fe^3+^ at 0 second (baseline) and 45 second (peak of the response); GFAP-eGFP images (green) are shown on the right. Scale bar, 10 μm. (C) Concentration-dependence of the maximal amplitude of [Ca^2+^]_i_ responses triggered by Fe^2+^ or Fe^3+^. The apparent EC_50_ are 0.635 μM for Fe^2+^ and 0.711 μM for Fe^3+^.

When imaging cortical astrocytes *in vivo* (the cells were identified by specific eGFP fluorescence) we found that addition of either Fe^2+^ or Fe^3+^ to the superfusing solution for 30 s induced transient [Ca^2+^]_i_ increase (Fig 1B). Administration of Fe^2+^ increased fluorescent intensity of Rhod-2 to 456.30% ± 18.46% (n = 10, p < 0.0001) whereas Fe^3+^ increased the peak of florescent signal to 308.50% ± 13.01% (n = 10, p < 0.0001) of the basal value.

### DMT1 and TFR mediate Fe^2+^ and Fe^3+^ uptake

As mentioned above, Fe^2+^ uptake is mediated by plasmalemmal transporter DMT1, whereas Fe^3+^ is accumulated in TF-bound form by TFRs (Fig. 2A). Immunostaining of cortical tissue preparations and primary cultured astrocytes demonstrated co-localisation of DMT1 and TFR with astroglial GFAP-positive profiles (Fig. 2B). In the cortical tissue both DMT1 and TFR showed preferential localisation at privascular endfeet. Meanwhile, expression of specific DMT1 and TFR mRNA was also detected in the freshly isolated and FACS-sorted astrocytes and neurones, as well as and in the cerebral tissues (Fig. 2C).

**Figure 2.**
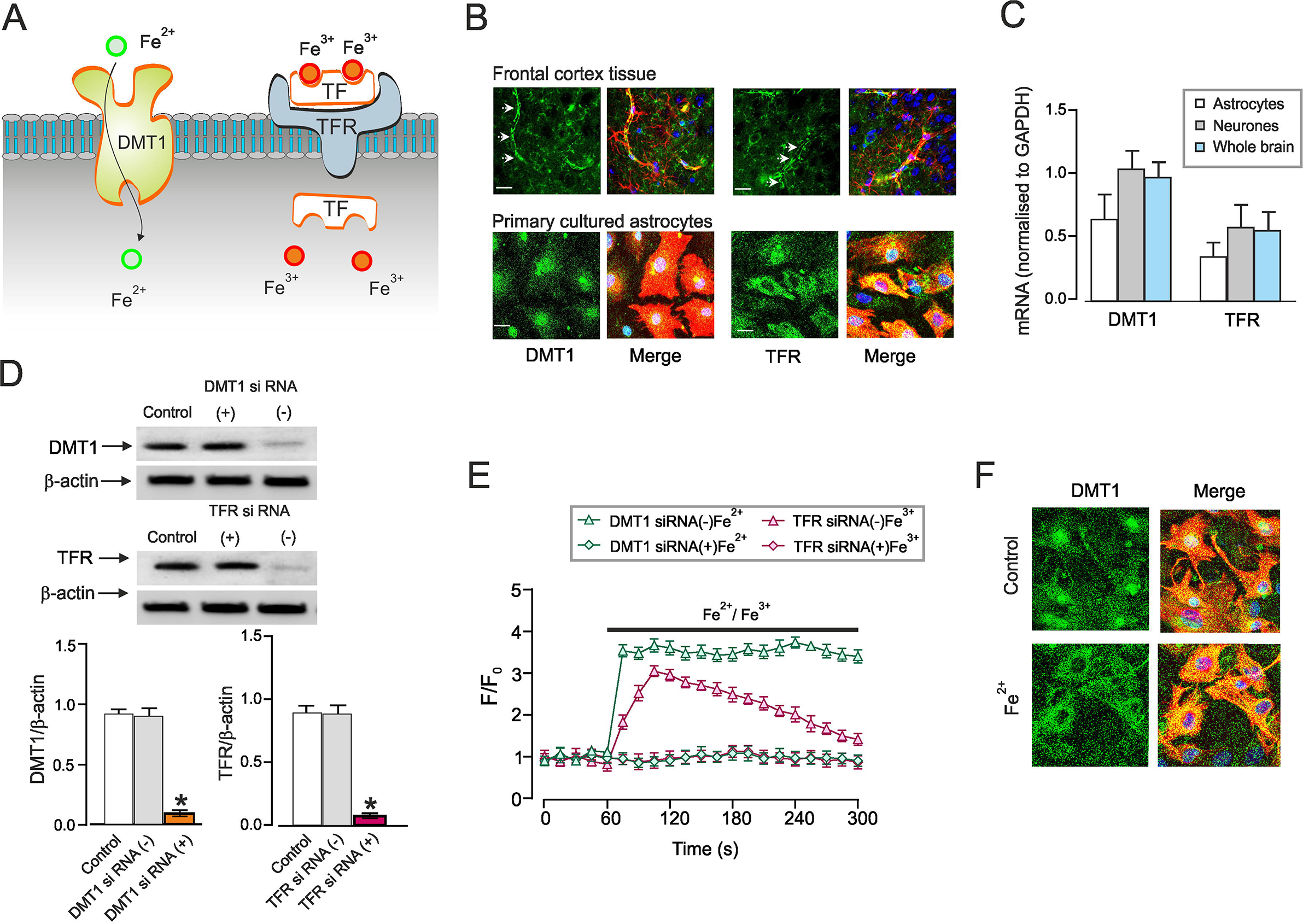
Pathways of Fe^2+^ or Fe^3+^ uptake. (A) Astrocytes accumulate Fe^2+^ through plasmalemmal divalent metal transporter 1 (DMT1/ SLC11A2) whereas Fe^3+^ is taken up by internalisation of Fe^3+^-TF-TFR complex. (B) Images of astrocytes in the frontal cortex preparations or in primary culture double-immunolabelled with antibodies against DMT1 or TFR and against GFAP. Scale bar, 10 μm. (C) The mRNA expression of DMT1 and TFR measured by qPCR in astrocytes sorted from GFAP-GFP mice, in neurones sorted from Thy1-YFP mice in whole cerebral tissues of wild type mice. (D) Representative western blot bands for DMT1 and TFR in cultured astrocytes treated with sham (Control), siRNA negative control (−) or positive duplex chains (+). The protein levels are shown as the ratio of DMT1 (55 kDa) and β-actin (42 kDa), and TFR (92 kDa) and β-actin. Data represent mean ± SD, n=6. *Indicates statistically significant (p<0.05) difference from the value of baseline in the same group. (E) [Ca^2+^]_i_ induced by Fe^2+^ or Fe^3+^ after RNA interference and down-regulation of protein synthesis. After treatment with DMT1 or TFR siRNA negative control (−) or positive duplex chains (+) for 3 days, [Ca^2+^]_i_ dynamics in response to Fe^2+^/Fe^3+^ was monitored. Every data point represents mean ± SD, n = 10. (F) Images of astrocytes in primary culture treated with sham (Control) or Fe^2+^ for 5 min and double-immunolabelled with antibodies against DMT1 and against GFAP. Scale bar, 10 μm.

To reveal the contribution of DMT1 and TFR to Fe^2+^/Fe^3+^-induced [Ca^2+^]_i_ dynamics, we inhibited expression of DMT1 or TFR using siRNA duplex chains. The representative protein western blots demonstrating the efficacy of knockdown are shown in Fig. 2D. When compared to the control group, the DMT1 siRNA reduced expression of DMT1 to 9.52% ± 2.58% (n = 6, p < 0.0001), whereas treatment with TFR siRNA decreased TFR levels to 7.84% ± 2.10% (n = 6, p < 0.0001). Administration of Fe^2+^ to DMT1-deficient astrocytes failed to induce any changes in [Ca^2+^]_i_. At the same time Fe^2+^ induced robust [Ca^2+^]_i_ elevation in astrocytes treated with siRNA (−) (Fig. 2E). Similarly, Fe^3+^ did not produce [Ca^2+^]_i_, transients in astrocytes treated with TFR siRNA duplex chains, whereas in cells exposed to negative control siRNA Fe^3+^ evoked [Ca^2+^]_i_ elevations (Fig. 2E). In un-stimulated astrocytes in culture the DMT1 fluorescence was the highest around the nucleus, suggesting its preferred intracellular localisation. After treatment of the cultures with Fe^2+^ for 5 minutes we observed redistribution of DMT1 from the nuclear region to the plasma membrane (Fig. 2F).

### Sources of iron-induced [Ca^2+^]_i_ mobilisation

The main sources of [Ca^2+^]_i_, increase in astrocytes are (i) Ca^2+^ release from the ER following opening of InsP_3_Rs or RyRs, or (ii) plasmalemmal Ca^2+^ influx through either Ca^2+^ permeable channels (such as L-type Ca^2+^ channels or TRP channels) or NCX operating in the reverse mode (Fig. 3A) or (iii) combination of some or all of these pathways. To dissect Ca^2+^ sources we first determined the influence of extracellular Ca^2+^ on iron-evoked [Ca^2+^]_i_ transients. Removal of Ca^2+^ from the extracellular milieu completely abolished Fe^2+^-induced [Ca^2+^]_i_ elevations but left Fe^3+^-evoked [Ca^2+^]_i_ transients largely intact (Fig. 3B). This highlighted the role for plasmalemmal Ca^2+^ influx in Ca^2+^ signalling triggered by Fe^2+^ and ER Ca^2+^ release for Ca^2+^ signals triggered by Fe^3+^.

**Figure 3.**
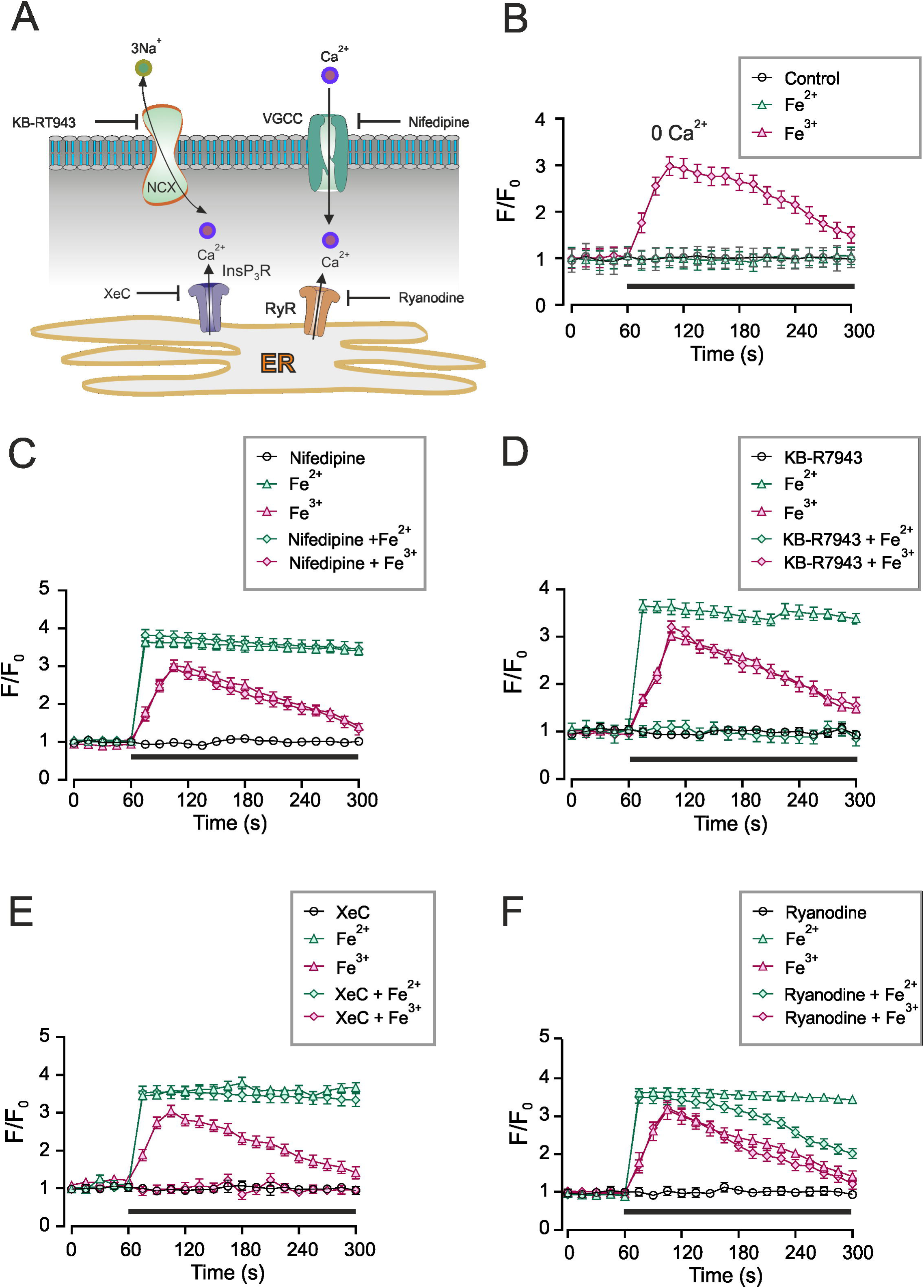
Ca^2+^ sources for Fe^2+^ or Fe^3+^-induced [Ca^2+^]_i_ transients. (A) Possible pathways mediating Ca^2+^ influx into the cytosol. (B) [Ca^2+^]_i_ recordings performed in the Ca^2+^-free extracellular medium; in the absence of Ca^2+^ exposure to F^e2+^ does not affect [Ca^2+^]_i_; conversely [Ca^2+^]_i_ transient evoked by Fe^3+^ remains intact. (C) Pre-treatment with 10 μM nifedipine does not affect Fe^2+^ or Fe^3+^ induced [Ca^2+^]_i_ responses. (D) Pre-treatment with NCX inhibitor KB-R7943 at 10 μM suppresses [Ca^2+^]_i_ response to Fe^2+^ but not to Fe^3+^. (E) Pre-treatment with InsP_3_ receptor inhibitor XeC at 10 μM suppresses [Ca^2+^]_i_ response to Fe^3+^ but not to Fe^2+^. (F) Pre-treatment with 10 μM ryanodine (Ry), the inhibitor of RyRs, affects only plateau phase of Fe^2+^-induced [Ca^2+^]_i_ response. In B – F every data point represents mean ± SD, n=10.

Incubation of astrocytes with an inhibitor of L-type voltage-gated Ca^2+^ channel nifedipine (10 μM) did not affect [Ca^2+^]_i_ responses to Fe^2+^ or to Fe^3+^ (Fig. 3C). In contrast, inhibition of NCX with selective agonist KB-R7943 at 10 μM completely eliminated [Ca^2+^]_i_ response to Fe^2+^, without affecting Fe^3+^-induced [Ca^2+^]_i_ transients (Fig. 3D). Thus, two forms of iron, the ferrous and ferric, mobilise intracellular Ca^2+^ through distinct pathways: Fe^2+^ stimulates Ca^2+^ influx by NCX, whereas Fe^3+^ triggers intracellular Ca^2+^ release. This suggestion was further corroborated by pharmacological inhibition of InsP_3_ receptors with potent antagonist Xestospongin C ^34^, Exposure of cultured astrocytes to 10 μM of XeC effectively suppressed [Ca^2+^]_i_ response to Fe^3+^, without much affecting Fe^2+^-induced [Ca^2+^]_i_ transient (Fig. 3E). Finally, treatment with 10 μM ryanodine (which at this concentration inhibits Ca^2+^-induced Ca^2+^ release ER channels) somewhat decreased the plateau phase of Fe^2+^ -induced [Ca^2+^]_i_ transient without modifying [Ca^2+^]_i_ response to Fe^3+^ (Fig. 3F).

### DMT1 transports Fe^2+^, which inhibits NKA, increases [Na^+^]_i_ and reverses NCX

Experiments described above have demonstrated that Fe^2+^, after being transported into the cell by DMT1, leads to a reversal of the NCX, which results in Ca^2+^ influx. The NCX reversal in astrocytes is usually triggered following an increase in the [Na^+^]_i_. Such an increase may originate either from the activation of plasmalemmal Na^+^ entry or from inhibition of the NKA, which maintains basal [Na^+^]_i_ ^19, 21^. The activity of NKA was acutely suppressed by exposure to 10 μM Fe^2+^ to 82.40 ± 5.74% (n = 10, p < 0.0001) of the control. Exposure to 100 nM of the specific NKA inhibitor, ouabain reduced NKA activity to 72.30 ± 5.91% (n = 10, p < 0.0001) of the control (Fig. 4A). When 10 μM Fe^2+^ and 100 nM ouabain were added together, the NKA activity was reduced further to 71.80 ± 7.81% (n = 10, p < 0.0001) (Fig. 4A).

**Figure 4.**
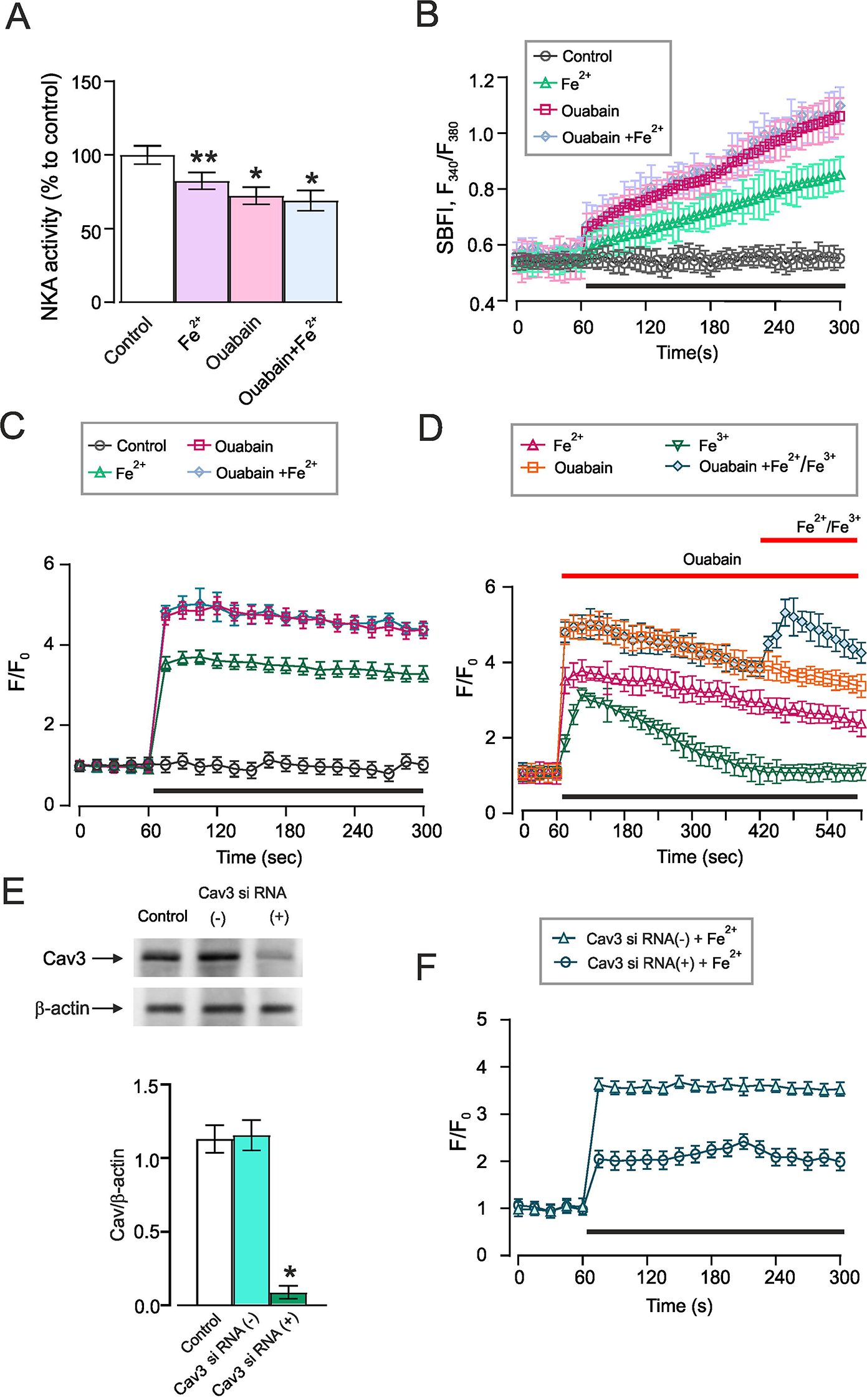
Fe^2+^ reverses NCXs by inhibiting NKA. (A) NKA activity in the presence of Fe^2+^ and ouabain measured by ELISA; OD value of was normalized to controls. Data represent mean ± SD, n=10. *Indicates statistically significant (p<0.05) difference from control group; **indicates statistically significant (p<0.05) difference from any other group. (B) [Na^+^]_i_ or (C) [Ca^2+^]_i_ changes induced by Fe^2+^ and ouabain. Every data point represents mean ± SD, n=10. (D) [Ca^2+^]_i_ recordings from cultured astrocytes challenged with Fe^2+^, Fe^3+^ and ouabain as indicated on the graph. (E) Representative western blot bands for Cav3 in cultured astrocytes treated with sham (Control), siRNA negative control (−) or positive duplex chains (+) to down-regulate Cav3 expression. The protein values are shown as the ratio of Cav3 (22 kDa) and β-actin (42 kDa). Data represent mean ± SD, n=6. *Indicates statistically significant (p<0.05) difference from any other group. F) [Ca^2+^]_i_ induced by Fe^2+^ after Cav3 RNA interference suppresses Fe^2+^-induced [Ca^2+^]_i_ response. Every data point represents mean ± SD, n=10.

Inhibition of NKA in astrocytes results in a substantial elevation in [Na^+^]_i_. When monitoring the [Na^+^]_i_ in cultured astrocytes with Na^+^-sensitive probe SBFI we found that both Fe^2+^ (10 μM) and ouabain (100 nM) triggered rapid and substantial elevation of [Na^+^]_i_ (Fig. 4B). When the cells were pre-treated with ouabain for 30 min, application of Fe^2+^ had no effect on [Na^+^]_i_ (n = 10, data not shown), whereas acute application of ouabain and Fe^2+^ lead to an increase in [Na^+^]_i_ (Fig 4B). These changes in [Na^+^]_i_ were paralleled by [Ca^2+^]_i_ dynamics. Exposure of astrocytes to Fe^2+^, ouabain or mixture of Fe^2+^ and ouabain caused [Ca^2+^]_i_ elevation (Fig. 4C). When Fe^2+^ was applied in the presence of ouabain it failed to change [Ca^2+^]_i_ (Fig. 4D); at the same time application of Fe^3+^ in the presence of ouabain still triggered additional [Ca^2+^]_i_ elevation (Fig. 4D).

### Fe^2+^-induced Ca^2+^ mobilisation is associated with caveolae

Treatment of cultured astrocytes with interfering Cav3 siRNA duplex chains decreased the level of Cav3 was decreased to 7.41 ± 4.32% (n = 6, p < 0.0001) of the control (Fig. 4E). An *in vitro* knock-down of Cav3 significantly reduced the man amplitudes of Fe^2+^-induced [Ca^2+^]_i_ increase; maximal increase in Fluo-4 F/F_0_ after Cav3 knockdown was 242.00 ± 16.44% (n = 10, p < 0.0001); whereas in control astrocytes treated with negative siRNA the amplitude of Fe^2+^-induced [Ca^2+^]_i_ increase reached 363.34 ± 11.62% (n = 10, p < 0.0001, Fig. 4F). To further analyse the effects of Cav3 on Fe^2+^-induced [Ca^2+^]_i_ dynamics the levels of relevant proteins were measured in the extracted caveolae (Fig. 5A). As shown in Fig. 5B, exposure to 10 μM Fe^2+^ for 5 min significantly increased the level of DMT1 to 293.24 ± 24.89% (n = 10, p < 0.0001) of the control values. After pre-treatment with Cav3 siRNA duplex chains, Fe^2+^ increased the level of DMT1 to only 195.77 ± 20.19% (n = 10, p < 0.0001) of control group (Fig. 5B). The levels of NCX1 and NKA were similarly affected by Fe^2+^ and the knocking down of Cav3. Exposure to Fe^2+^ increased the level of NCX1 and NKA to 248.71 ± 19.58% (n = 10, p < 0.0001) and 263.66 ± 25.93% (n = 10, p < 0.0001) of the controls, after knock down Cav3, Fe^2+^ only elevated the level of NCX1 and NKA to 172.96 ± 11.76% (n = 10, p < 0.0001) and 200.86 ± 18.14% (n = 10, p < 0.0001) of control values (Fig. 5B). Of note, Fe^2+^ did not affect the levels of NCX2 and NCX3 (Fig. 5B).

**Figure 5.**
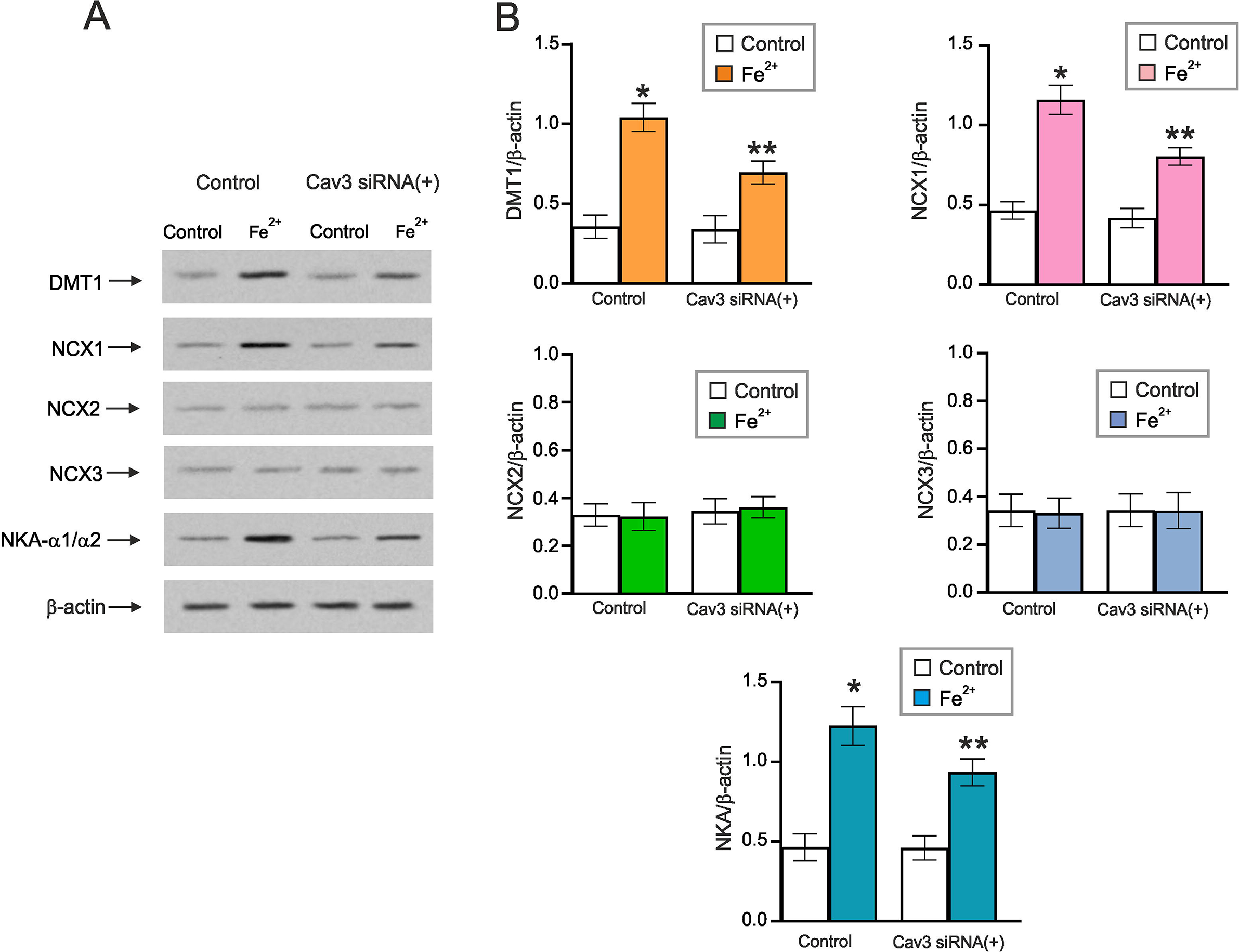
Caveolae integrate DMT1, NKA and NCX. (A) Representative protein western blot bands for DMT1, NCX1-3 and NKA-α1/α2 in caveolae membranes extracted from cultured astrocytes. Astrocytes were treated with sham (Control), with siRNA negative control (−) or with Cav3 duplex chains (siRNA(+)) for 3 days. (B) Protein levels are shown as the ratio of DMT1 (55 kDa) and β-actin (42 kDa), NCX1 (108 kDa) and β-actin, NCX2 (102 kDa) and β-actin, NCX3 (100 kDa) and β-actin, NKA-α1/α2 (100 kDa) and β-actin (42 kDa). Data represent mean ± SD, n=6. *Indicates statistically significant (p<0.05) difference from any other group; **indicates statistically significant (p<0.05) difference from control plus Fe^2+^ group or Cav3 siRNA(+) plus Ctrl group.

### Fe^3+^ triggers Ca^2+^ release through stimulation of PLC and increase in InsP_3_ production

As shown in Fig. 3E inhibition of InsP_3_ receptors with XeC suppressed Fe^3+^-induced [Ca^2+^]_i_ mobilisation. We therefore analysed affects of Fe^3+^ on the InsP_3_ signalling cascade in cultured astrocytes. Incubation of astrocytes with Fe^3+^ increased the level of InsP_3_ in cell lysates to 74.10 ± 8.14 ng/ml (n = 10, p < 0.0001), from the resting InsP_3_ level of 24.40 ± 6.35 ng/ml in control group. In cells treated with TF alone the InsP_3_ level was 28.00 ± 12.62 ng/ml (n = 10, p = 0.4309) (Fig. 6A). Subsequently we analysed the links between scaffolding/signalling protein Dab2 and Fe^3+^-induced Ca^2+^ signalling. We suppressed expression of two isoforms of Dab2 by siRNA duplex chains (Fig. 6B). After RNA interfering, expressions of 96KD and 67KD Dab2 isoforms decreased, respectively, to 5.81 ± 3.56% (n = 6, p < 0.0001) and to 11.88 ± 6.51% (n = 6, p < 0.0001) of the control values (Fig. 6B). The knock-down of Dab2 rendered Fe^3+^ ineffective: exposure to Fe^3+^ in Dab2-deficient astrocytes did not affect InsP_3_ production (control: 31.60 ± 12.51 ng/ml (n = 10); Dab2 knockdown: 30.60 ± 10.42 ng/ml (n = 10, p = 0.1253; Fig. 6C).

**Figure 6.**
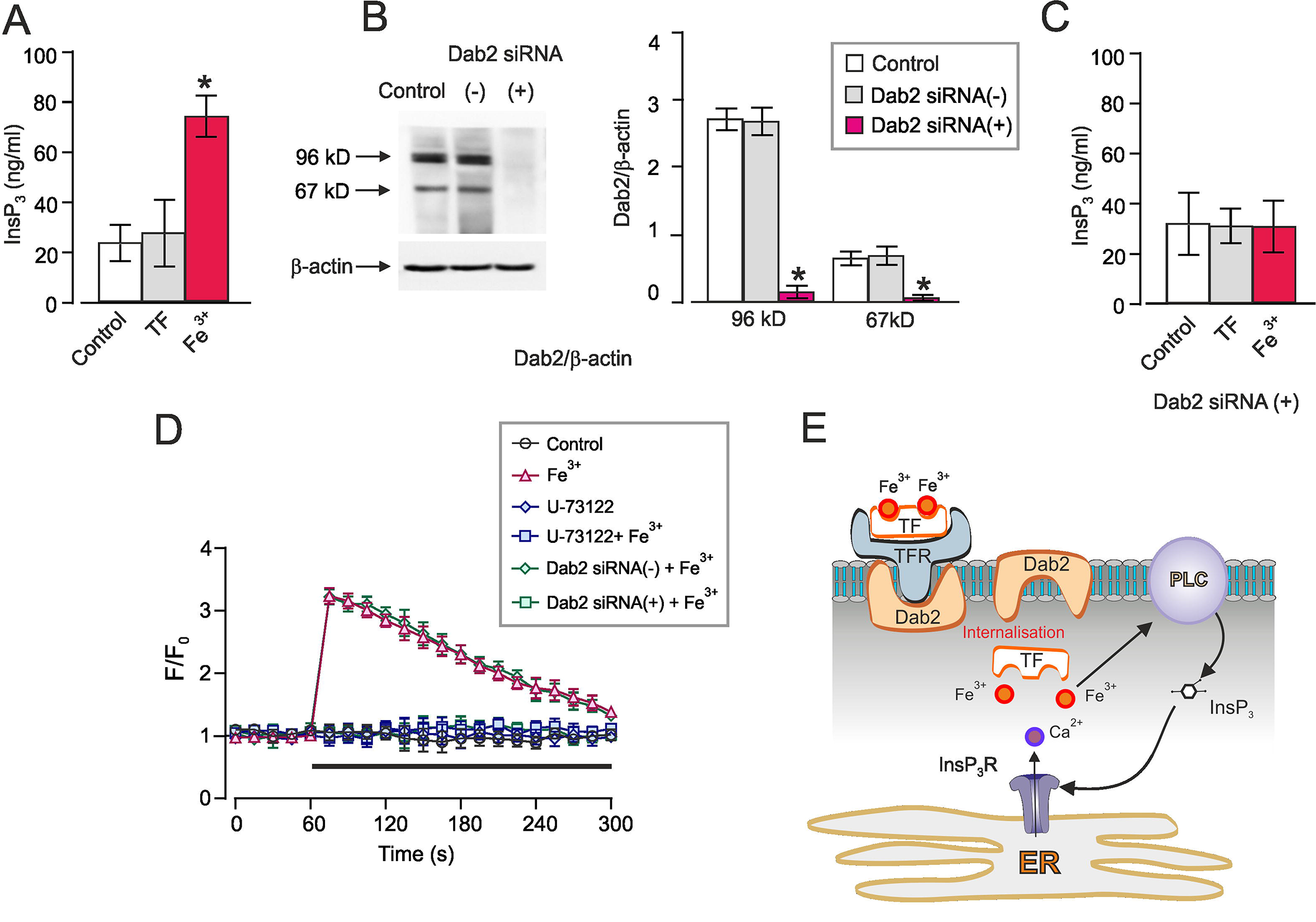
Mechanisms of Fe^3+^ induced Ca^2+^ signalling. (A) Incubation with Fe^3+^ increases InsP_3_ level in cultured astrocytes; InsP_3_ was measured with ELISA; data are presented as mean ± SD, n=10. *Indicates statistically significant (p<0.05) difference from any other group. (B) Representative protein bands two isoforms (96KDa and 67KDa) of Dab2 in cultured astrocytes treated with sham (Control), siRNA negative control (−) or positive duplex chains (+). The protein values are shown as the ratio of 96 kDa isoform and β-actin (42 kDa), and the ratio of 67 kDa isoform and β-actin. Data represent mean ± SD, n=6. *Indicates statistically significant (p<0.05) difference from any other group. (C) Fe^3+^-dependent changes in the InsP_3_ after treatment with Dab2 siRNA duplex chains. After treatment with Dab2 siRNA duplex chains (+) for 3 days, the primary cultured astrocytes were treated with serum-free medium (Control), TF (as negative control) or 10 μM Fe^3+^-TF (Fe^3+^) for 5 minutes, the level of InsP_3_ was measured by ELISA and shown as mean ± SD, n=10. (D) Fe^3+^ -induced [Ca^2+^]_i_ responses are Dab2 and PLC dependent. Treatment with Dab2 siRNA duplex chains (+) as well as with 10 μM U-73122 (PLC inhibitor) suppressed Fe^3+^-induced [Ca^2+^]_i_ responses. Every data point represents mean ± SD, n=10. (E) Mechanisms of Fe^3+^-induced Ca^2+^ signalling in astrocytes. The uptake of Fe^3+^ is mediated TFR, TFR internalization requires Dab2; when in the cytosol Fe^3+^ activated PLC thus stimulating InsP3-induced Ca^2+^ release from the ER.

When Dab2-deficient astrocytes were challenged with Fe^3+^, no [Ca^2+^]_i_ increase was recorded (Fig. 6D). Similarly, after inhibition of the PLC with U-73122, application of Fe^3+^ did not change [Ca^2+^]_i_ (Fig. 6D). Hence, we may surmise that uptake of Fe^3+^ through TFR requires Dab2 protein; after entering the cytosol Fe^3+^ activates the PLC, which produces InsP_3_ that triggers InsP_3_-induced Ca^2+^ release from the ER (Fig. 6E).

### Iron increases phosphorylation of cPLA2 and stimulates secretion of arachidonic acid and prostaglandin E2

To characterise downstream cascades induced by iron ions we measured phosphorylation of cPLA2 and NF-κB. As compared to the control group, Fe^2+^ and Fe^3+^ increased the ratio of p-cPLA2 and cPLA2 respectively to 210.00 ± 19.38% (n = 10, p < 0.0001) and 177.42 ± 15.05% (n = 10, p < 0.0001) (Fig. 7A, B). Similarly, the ratio of the phosphorylated P65 and P65 was increased by Fe^2+^ and Fe^3+^ to 164.58 ± 10.66 (n = 10, p < 0.0001) and 141.20 ± 11.85% (n = 10, p < 0.0001) of the control values (Fig. 7A, C). The pre-treatments with Ca^2+^ chelator BAPTA-AM abolished an increase in the phosphorylation of cPLA2 following exposure to Fe^2+^ or Fe^3+^ to 100.65 ± 19.93% (n = 10, p = 0.9429) and 83.87 ± 17.86% (n = 10, p = 0.0716) of the control. Likewise, BAPTA-AM suppressed phosphorylation of P65 stimulated by Fe^2+^ or Fe^3+^ to 90.28 ± 13.80% (n = 10, p = 0.0939) and 96.99 ± 19.26% (n = 10, p = 0.6700) of the controls (Fig. 7A, C). Treatment with NCX inhibitor KB-R7943 and InsP_3_ receptor blocker Xe-C also abolished the phosphorylation of cPLA2 and P65 (Fig. 7A-C). The KB-R7943 decreased the phosphorylation of cPLA2 induced by Fe^2+^ to 98.46 ± 20.20% (n = 10, p = 0.8534) of control group, while Xe-C suppressed the Fe^3+^-stimulated phosphorylation of cPLA2 to 103.75 ± 23.32% (n = 10, p = 0.7065) of the control value. These two blockers had the same effect on phosphorylation of P65: Xe-C decreased the Fe^2+^-induced phosphorylation of P65 to 99.51 ± 16.82% (n = 10, p = 0.9418) of the control, whereas Xe-C decreased phosphorylation of P65 stimulated Fe^3+^ to 102.86 ± 17.71% (n = 10, p = 0.7048) of the control (Fig. 7B, C).

**Figure 7.**
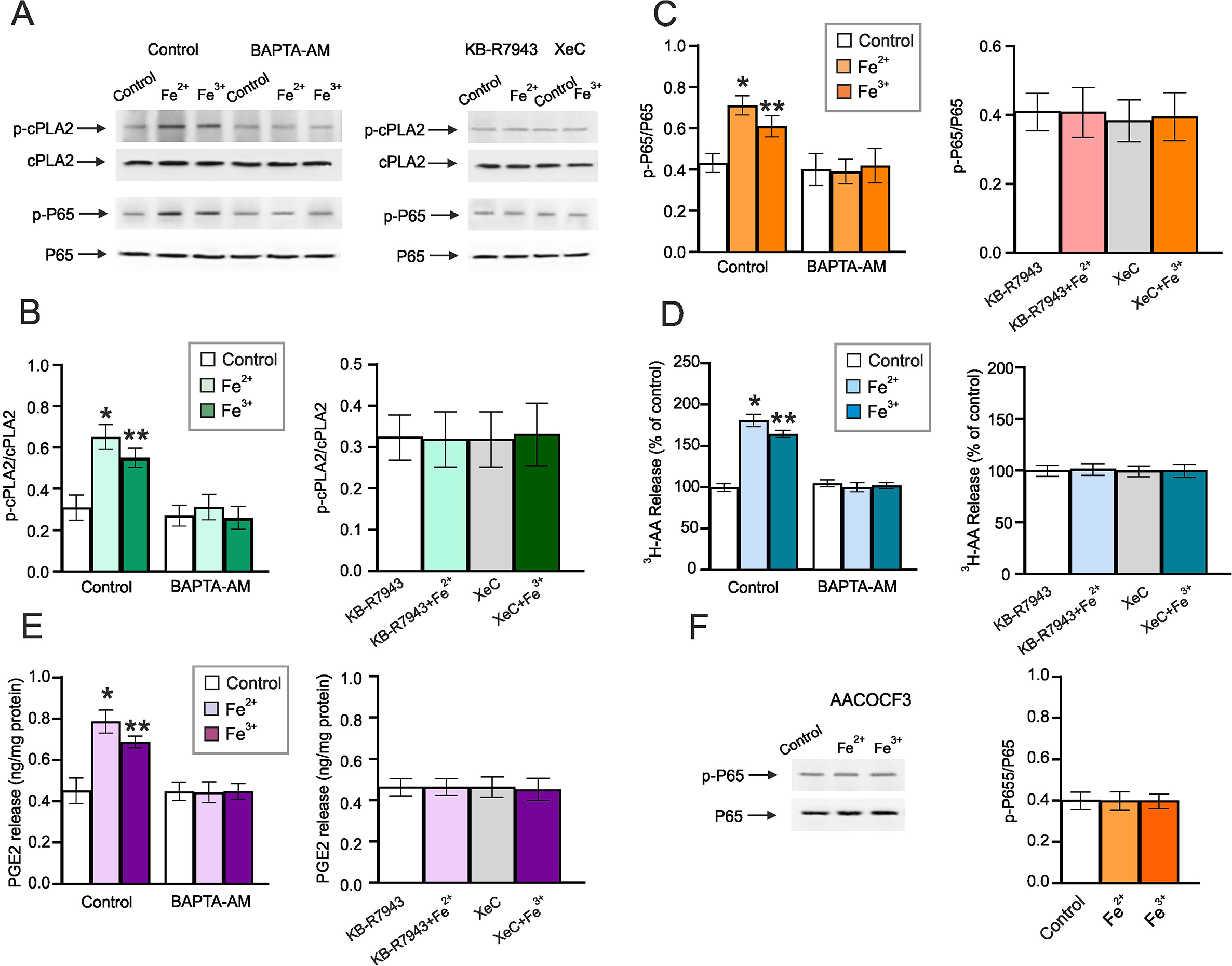
Fe^2+^ and Fe^3+^ induced Ca^2+^ signals stimulate phosphorylation of cPLA2 and NF-κB. (A - C) Representative protein bands for p-cPLA2, cPLA2, p-P65 and P65 subunit of NF-κB in cultured astrocytes. After pre-treatment with serum-free medium (Control) or 1 μM BAPT-AM (a Ca^2+^ chelator), or 10 μM KB-R7943 or 10 μM Xe-C for 30 minutes, astrocytes were treated with serum-free medium (Control), 10 μM Fe^2+^ or 10 μM Fe^3+^ for 5 minutes. Protein levels are shown as ratio of p-cPLA2 and cPLA2 (85 kDa) (B) and the ratio of p-P65 and P65 (65 kDa) (C). (D and E) After pre-treatment with serum-free medium (Control) or 1 μM BAPT-AM, 10 μM KB-R7943 or 10 μM Xe-C for 30 minutes, astrocytes were treated with serum-free medium (Control), 10 μM Fe^2+^ or 10 μM Fe^3+^-TF (Fe^3+^) for 5 minutes. Sectered levels of AA (D) and PGE2 (E) are shown normalized to control values. Data represent mean ± SD, n=10. *Indicates statistically significant (p<0.05) difference from any other group (B-E); **indicates statistically significant (p<0.05) difference from control + Fe^2+^ group (B-E). (F) Representative Western blot bands of p-P65 and P65 subunit of NF-κB in cultured astrocytes. After pre-treated with 2 μM AACOCF3 for 30 minutes, astrocytes were treated with serum-free medium (Control), 10 μM Fe^2+^ or Fe^3+^-TF (Fe^3+^) for 5 minutes. The protein levels are shown as the ratio of p-P65 and P65. Data represent mean ± SD, n=10.

We also measured secretion of AA and PGE2 using ELISA. As compared with control group, Fe^2+^ and Fe^3+^ increased extracellular level of AA to 180.90 ± 7.43% (n = 10, p < 0.0001) and 164.60 ± 4.35% (n = 10, p < 0.0001) of the control. Pre-treatment with BAPTA-AM (that eliminates any increase in [Ca^2+^]_i_) completely inhibited Fe^2+^ and Fe^3+^ -induced rise in AA secretion (to 100.10 ± 5.43% (n = 10, p = 0.9651) and 102.00 ± 3.74% (n = 10, p = 0.3014) of the controls (Fig. 7D). Similarly, inhibitors of NCX and InsP_3_ receptors eliminated effects of Fe^2+^ and Fe^3+^. Incubation with KB-R7943 decreased the level of AA in the presence of Fe^2+^ to 101.70 ± 5.06% (n = 10, p = 0.5803) of the control, while XeC suppressed the level of AA induced by Fe^3+^ to 100.40 ± 5.70% (n = 10, p = 0.8951) of the control (Fig. 7D). Treatment with Fe^2+^ and Fe^3+^ also stimulated secretion of PGE2; the levels of the latter increased, respectively, to 174.12 ± 12.39% (n = 10, p < 0.0001) and 152.21 ± 6.24% (n = 10, p < 0.0001) of the control values. Again, pre-treatment with BAPTA-AM completely inhibited stimulatory effects of iron on secretion of PGE2 (levels of PGE2 in the presence of Fe^2+^ and Fe^3+^ were 98.23 ± 11.19% (n = 10, p = 0.7538) and 99.33 ± 8.44% (n = 10, p = 0.8968) of the controls (Fig. 7E). Likewise, KB-R7943 suppressed Fe^2+^-induced, whereas XeC inhibited Fe^3+^-indueced stimulation of PGE2 release (with PGE2 values of 99.79 ± 8.48% (n = 10, p = 0.9550) 97.42 ± 11.38% (n = 10, p = 0.5991) of the controls (Fig. 7E). Finally, Fe^2+^ and Fe^3+^-induced phosphorylation of P65 was blocked by the specific inhibitor of cPLA2, AACOCF3 (Fig. 7F). After the pre-treatment with 2 μM AACOCF3, the level of p-P65 in the presence of Fe^2+^ was 98.77 ± 10.48% (n = 10, p = 0.7795) and the level of p-P65 in the presence of Fe^3+^ was 98.77 ± 7.81% (n = 10, p = 0.7451).

## Discussion

In this paper, we describe previously unknown effects of iron ions on cellular [Ca^2+^]_i_ in astrocytes. Administration of either Fe^2+^ or Fe^3+^ triggered a concentration-dependent increase in [Ca^2+^]_i_ with EC_50_ of 0.635 μM for Fe^2+^ and 0.711 μM for Fe^3+^. We further performed an in depth analysis of the mechanisms underlying iron transport and iron induced Ca^2+^ signalling. We also demonstrated that, contrary to the previous beliefs, astrocytes express functional TFR *in vitro* and *in vivo* thus allowing accumulation of Fe^3+^.

### Iron transport in astrocytes is mediated by DMT1 and TFR

Glial cells, and astrocytes in particular, store up to 75% of ionised iron in the CNS ^35^, being arguably active players in brain protection against iron overloads ^36^. Transmembrane transport of iron in astroglial cells is has not been studied in details. There is a general agreement of primary role of plasmalemmal divalent metal transporter 1, DMT1/SLC11A2, which selectively transports Fe^2+^; the DMT1 was identified in astrocytes in culture and there are limited data indicating its presence in astroglial endfeet *in situ* ^37–39^. The Fe^2+^ was also suggested to enter reactive astrocytes by diffusion through transient receptor potential “canonical” (TRPC) channels ^36^. Expression of TF-Fe^3+^-transporting TFR has been noted in astrocytes in culture (ref ^12, 40^); it is, however, generally believed that astrocytes *in vivo* are not in a possession of TFR and hence can not accumulate Fe^3+^ ^41–43^. This conclusion, however, has been made on the basis of rather limited investigations ^40, 44^; and expression of TFR-specific mRNA was detected in astroglial transriptome ^45^. In our study we confirmed expression of DMT1, at mRNA and protein levels as well as by immunostaining, in acutely isolated astrocytes, in astroglial primary culture and *in situ* in cortical tissue; the DMT1 was particularly enriched in the endfeet (Fig. 2B-D). Subsequently we detected astroglial TFR expression at mRNA level in the transcriptase of astutely isolated and FACS-sorted astrocytes (Fig. 2C). We further confirmed expression of TFR in astrocytes at a protein level and in immunohistochemical analysis of astrocytes in culture and in cortical preparations (Fig. 2B-D). In the the cortical tissue TFR labelling was concentrated in perivascular astrocytic endfeet (Fig. 2B).

### Mechanisms of iron induced Ca^2+^ signalling

Not much is known about the links between ionised iron and Ca^2+^ signalling in the cellular elements of the CNS. In the literature, we found only a single example of Fe^3+^-induced [Ca^2+^]_i_ transient in cultured hippocampal neurones ^46^. To the best of our knowledge here we present the first recordings of Fe^2+^/Fe^3+^-induced Ca^2+^ signals in astrocytes. Both ions evoked [Ca^2+^]_i_ elevation in primary cultured astrocytes and when administered to the cortices of alive animals studied with transcranial confocal microscopy. Both ions acted in the low μM range of concentrations, however the kinetics of [Ca^2+^]_i_ transients are different. Exposure of cultured astrocytes to Fe^2+^ triggered rapid [Ca^2+^]_i_ increase with long-lasting plateau; the [Ca^2+^]_i_ barely declined in the presence of Fe^2+^. In contrast, Fe^3+^-induced transient elevation of [Ca^2+^]_i_ recovered to the baseline within ~ 200 – 300 s in the presence of Fe^3+^ (Fig. 1B, 2E). These distinct kinetics reflect different signalling cascades activated by iron ions.

The Fe^2+^-induced [Ca^2+^]_i_ responses require DMT1; *in vitro* knockdown of DMT1 expression with silencing mRNA completely eliminated Ca^2+^ signal (Fig. 2). The Fe^2+^-induced [Ca^2+^]_i_ changes originate from plasmalemmal Ca^2+^ entry, because removal of Ca^2+^ from the extracellular medium inhibited [Ca^2+^]_i_ response. Finally, Fe^2+^-induced [Ca^2+^]_i_ signals require NCX, as pharmacological inhibition of the latter effectively suppressed [Ca^2+^]_i_ elevation (Fig. 3). These data indicate that Fe^2+^, after being accumulated in the astrocyte, switches the NCX into the reverse mode of operation which generates Ca^2+^ influx into the cell in exchange for Na^+^. This scenario requires increase in astroglial [Na^+^]_i_, which readily reverses the NCX ^19, 47^. An increase in [Na^+^]_i_ is likely to follow an inhibition of NKA, which represents the major Na^+^ efflux mechanism in astroglial cells ^21^. Activity of NKA indeed was suppressed by Fe^2+^, and probing astrocytes with Na^+^-sensitive indicator SBFI revealed Fe^2+^-induced [Na^+^]_i_ elevation (Fig. 4). These effects of Fe^2+^ were eliminated by NKA inhibitor ouabain thus demonstrating the central role of NKA in Fe^2+^-induced Ca^2+^ signalling (Figs. 4, 8).

**Figure 8.**
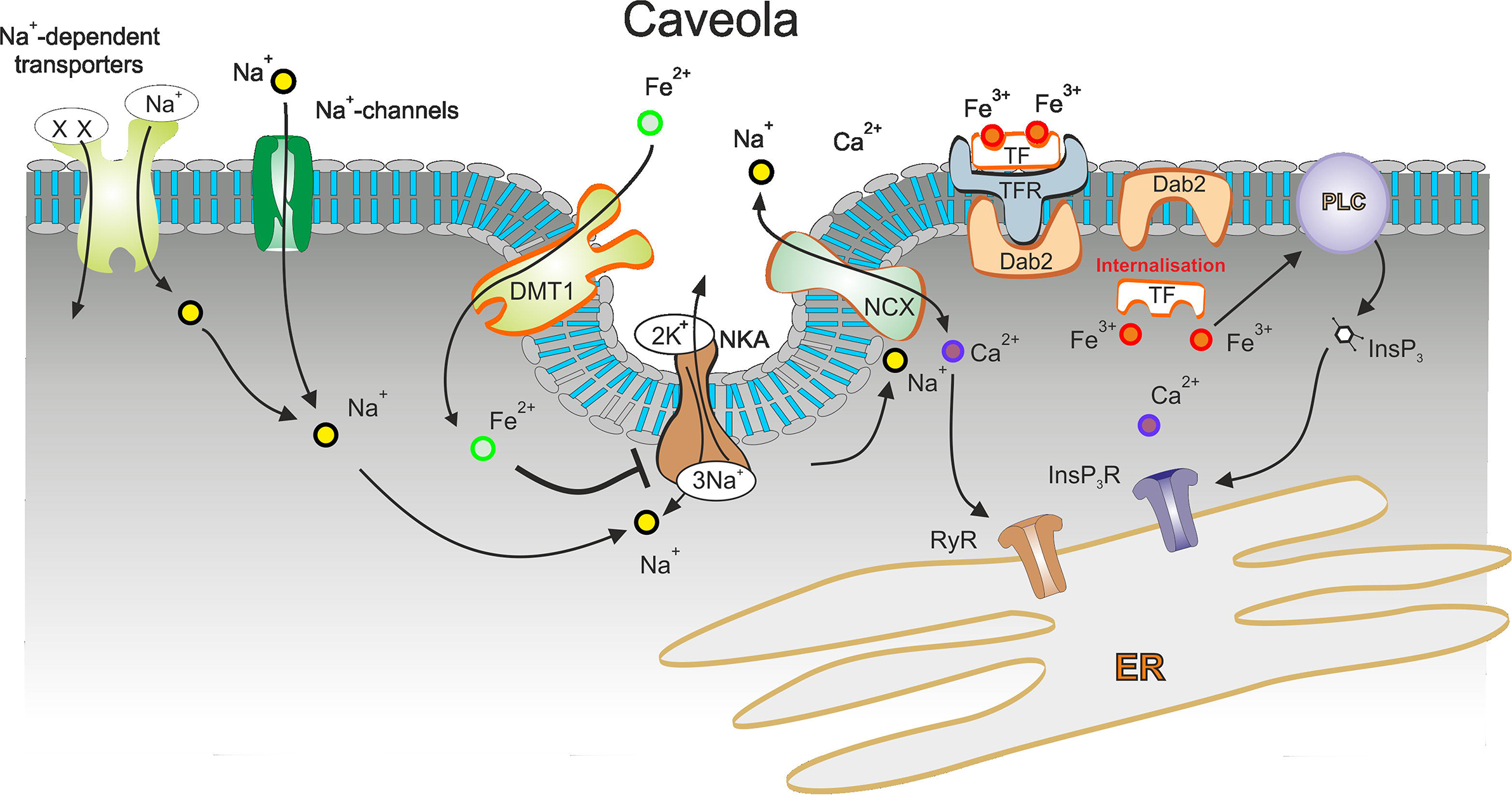
Mechanisms of Fe^2+^ and Fe^3+^ -induced intracellular Ca^2+^ signalling. Uptake of Fe^2+^ is mediated by DMT1 and uptake of Fe^3+^ is mediated by TFR. After entering the cytosol, Fe^2+^ inhibits the activity of NKA, which causes an increase in [Na^+^]_i_ due to an unopposed influx of Na^+^ through Na^+^-dependent transporters or Na^+^-channels. Increase in [Na^+^]_i_ switches the NCX into the reverse mode, which results in Ca^2+^ influx. This influx also activated Ca^2+^-induced Ca^2+^ release through RyR that contributes to the plateau phase of [Ca^2+^]_i_ response. Accumulation of Fe^2+^ also promotes the formation of the functional unit of DMT1, NCX1 and NKA in the caveolae by recruiting Cav3. Uptake of Fe^3+^ proceeds through Dab2-assisted internalization; after entering the cytosol Fe^3+^ activates the PLC and stimulates InsP_3_-dependent Ca^2+^ release from the ER.

The mechanism of Fe^3+^-induced Ca^2+^ signalling is associated with intracellular Ca^2+^ release. The Fe^3+^-induced [Ca^2+^]_i_ responses were preserved in Ca^2+^ free extracellular solution while being blocked by XeC (inhibitor of InsP_3_ receptor) and by U-73122 (inhibitor of PLC) thus revealing the central role for InsP_3_-medaited ER Ca^2+^ release. Initiation of this signalling cascade requires transmembrane transport of Fe^3+^; *in vitro* knockdown of TFR eliminated Fe^3+^-evoked [Ca^2+^]_i_ dynamics. The internalisation of TF-Fe^3+^-TFR complex also requires functional Dab2 protein. This protein is a multi-modular scaffold protein with signalling roles in ion homeostasis, inflammation and receptors internalization ^48^. For example, Dab2 influences signalling pathways of fibrinogen and its receptors (integrin α_IIb_β_3_) by regulating the complex internalization thus modulating the platelet aggregation ^48, 49^. In addition, Dab2 facilitates the binding of the internalized integrin α_IIb_β_3_ and phosphatidylinositol 4,5-bisphosphate (PIP2) in the activated platelet ^48, 50^. Ablation of Dab2 in astrocytes with specific siRNA interrupts signalling chain and blocks Fe^3+^-dependent Ca^2+^ signalling (Figs. 6, 8).

### Role of caveolae in iron-induced Ca^2+^ signalling

Caveoale are specific plasmalemmal structures that form functional microdomains involved in various signalling, endocytotic and transporting events ^51^. Caveolae and its main structural and regulatory proteins Caveolin-1,2,3 are present in astrocytes (with predominant expression of Cav3); astroglial caveolae contribute to signal transduction, formation of signalling protein complexes and are involved in action of various neuroactive substances and drugs ^52, 53^. Caveolae are known to form functional Ca^2+^ signalling units, establish links between Ca^2+^ channels and various transports and may be a substrate for plasmalemmal/ER functional domains operational in astrocytes ^54^. We found that down-regulation of expression of Cav3 in cultured astrocytes substantially reduced the amplitude of Fe^2+^-evoked [Ca^2+^]_i_ responses. We suggest therefore that Cav3 and caveolae integrate DMT1, NKA and NCX into a single Ca^2+^ signalling unit (Fig. 8) and moreover exposure to iron increases formation of such units.

### Do astrocytes protect the brain against iron overload?

Iron homeostasis is of fundamental importance for cells, tissues and organisms, as iron contributes to a wide range of vital biological pathways ^55^. The brain contains high concentrations of bound and free iron, which participates in multiple processes from energy production to synaptic transmission ^35^. Iron overload and failures in iron homoeostatic cascades triggers neurotoxicity and is implicated in brain diseases ^42^. Genetic mutation of iron regulatory proteins (the key elements of iron homoeostasis) results in iron deposition in the brain with subsequent neurodegeneration characteristic for aceruloplasminemia ^56^ and neuroferritinopathy ^57^. Similarly, iron accumulation has been characterised in neurodegenerative diseases including Alzheimer’s disease, Parkinson disease, amyotrophic lateral sclerosis and Huntington disease to name but a few ^42^. Specific class of neurodegeneration with brain iron accumulation (NBIA) has been also categorised in recent years ^58^.

Based on our data we propose that astrocytes mount the defence against iron overload. This defence includes iron accumulation through both DMT1 and TFR, redistribution of DMT1 from intracellular locations to plasmalemma and generation of Ca^2+^ signals, which stimulate NF-κB cascade and increase secretion of neuroprotective molecules such as arachidonic acid and prostaglandin E2. Iron-induced Ca^2+^ signalling is activated by low pathological iron concentrations (> 1 μM; while physiological iron concentration in the CSF ranges between 0.3 and 0.75 μM). Importantly, two distinct signalling cascades (DMT1 Fe^2+^ transport, inhibition of NKA and reversal of NCX versus Fe^3+^-TF-TFR transport, activation of PLC and generation of InsP_3_-induced Ca^2+^ release) distinguish between ferric and ferrous. These distinct pathways may define very different outputs: it is known for example that activation of astroglial InsP_3_ receptors type II is linked to initiation of reactive astrogliosis ^59, 60^. Astrogliosis plays important, if not defining, role in the evolution of many neurological diseases ^61^. Our previous experiments have shown that formation of brain deposits of iron up-regulates astroglial expression of TFR and instigates reactive astrogliosis ^62^. What characterises the iron-induced reactive phenotype and what is the role of astroglial reactivity in managing excessive iron in the brain remains to be found. In conclusion, our study presents a novel phenomenon that iron ions (Fe^2+^ and Fe^3+^) directly induce intracellular Ca^2+^ signalling and stimulate astroglial protective mechanisms against iron overload in broad pathological contexts.

## MATERIALS and METHODS

### Materials

The culture medium including DMEM and foetal bovine serum were purchased from Gibco Life Technology Invitrogen (Grand Island, NY, USA). Oligo-fectamine, MEMI, fluo-4 AM, sodium-binding benzofuran isophthalate (SBFI) AM, G-agarose bead, TFR antibody, β-actin antibody, GFAP antibody, DMT1 antibody and DMT1 siRNA duplex chains were from Thermo Fisher Scientific (Waltham, MA USA); siRNA duplex chains of TFR, NCX1-3, Cav-3 and Dab2, NCX2 antibody, p-cPLA2 antibody, cPLA2 antibody, p-NF-κB P65 antibody, NF-κB P65 antibody, Na^+^/K^+^-ATPase alpha1/2 antibody and the secondary antibodies were bought from Santa Cruz Biotechnology (Santa Cruz, CA, USA). NCX1 antibody, NCX3 antibody and native mouse apo-transferrin (apo-TF; i.e. iron-free) were from Abcam (Cambridge, MA, USA). Donkey serum, xestospongin C (Xe-C), nifedipine, AACOCF3, BAPTA-AM, ferrous sulfate heptahydrate (FeSO_4_) and ferric ammonium citrate were purchase from Sigma-Aldrich (St. Louis, MO, USA). Ryanodine and KB-R7943 was purchased from Calbiochem (La Jolla, CA, USA). Secondary antibody staining with donkey anti-mouse or anti-rabbit Cy-2/3 were from Jackson Immuno-Research (West Grove, PA, USA).

### Animals

The C57BL/6 mice, FVB/N-Tg(GFAP-eGFP)14Mes/J and B6.Cg-Tg(Thy1-YFP)HJrs/J transgenic mice were all purchased from the Jackson Laboratory (Bar Harbor, ME, USA). The animals were raised in standard housing conditions (22 ± 1◻; light/dark cycle of 12/12h), with water and food available *ad libitum*. All experiments were performed in accordance with the US National Institutes of Health Guide for the Care and Use of Laboratory Animals (NIH Publication No. 8023) and its 1978 revision, and all experimental protocols were approved by the Institutional Animal Care and Use Committee of China Medical University, No. [2019]059.

### Primary culture of astrocytes

Astrocytes were cultured from newborn mice as described previously ^63, 64^. In brief, the cerebral hemispheres were isolated, dissociated and filtered. Isolated astrocytes were grown in Dulbecco’s Minimum Essential Medium (DMEM) with 7.5 mM glucose supplemented with 10% foetal bovine serum. Astrocytes were incubated at 37 ◻ in a humidified atmosphere of CO_2_/air (5:95%). The cultures are highly enriched in astrocytes, the purity is >95% as judged by GFAP staining ^65^.

### Iron Treatments

For preparing Fe^3+^-TF solution, ferric ammonium citrate and mouse apo-TF were incubated at a 2:1 ratio in serum-free culture medium for 1 hour at 37 ◻ ^66, 67^. The same concentration of apo-TF in the same volume of culture medium but without Fe^3+^ was used for control treatments. For the Fe^2+^ solution, FeSO_4_ was freshly dissociated in serum-free culture medium at 37 ◻ and used immediately, the same volume of serum-free culture medium was used as the control for Fe^2+^ group.

### RNA Interfering

As described previously ^65, 68, 69^, cultured astrocytes were incubated in DMEM without serum for 12 hours before transfection. A transfection solution containing 2 μl oligo-fectamine (Promega, Madison, WI, USA), 40 μl MEMI, and 2.5 μl siRNA (DMT1, TFR, NCX1-3, Cav-3 or Dab2) was added to the culture medium in every well for 8 h. In the siRNA-negative control cultures, transfection solution without siRNA was added. Thereafter, DMEM with three times serum was added to the cultures. These siRNA duplex chains were purchased from Santa Cruz Biotechnology (CA, USA).

### Preparation of Membrane Caveolae

Cell homogenization and the caveolae preparation from astrocytes was made as previously described ^70, 71^. In brief, primary cultured astrocytes were collected and homogenised in SET (0.315 M sucrose, 20 mM TrisCl, and 1 mM EDTA, pH 7.4), and centrifuged for 1 h at 1,000×g. The pellets were re-solubilised in SET and layered on Percoll (30% in SET) followed centrifugation at 1,000×g. The pellets were re-homogenised and re-layered to three sucrose density gradient solution (80%, 30% and 5%) with ultra-centrifugation at 175,000×g. Finally, the purified caveolae were collected and re-suspended in SET.

### co-Immunoprecipitation

We used technologies of co-immunoprecipitation and subseqeunet western blotting to check the conjunction level between NCXs and DMT1, as described previously ^68^. After homogenization, protein content was determined by the Bradford method ^72^ using bovine serum albumin as the standard. For immunoprecipitation of NCX1-3, whole cell lysates (500 μg) were incubated with 20 μg of anti-NCX1, anti-NCX2 or anti-NCX3 antibody for overnight at 4 °C. Then, 200 μl of washed protein G-agarose bead slurry was added, and the mixture was incubated for another 2 hours at 4 °C. The agarose beads were washed three times with cold phosphate buffer solution (PBS) and collected by pulsed centrifugation (5 s in a microcentrifuge at 14,000×*g*), the supernatant was drained off, and the beads were boiled for 5 min. Thereafter, the supernatant was collected by pulsed centrifugation, and the entire immunoprecipitates were subjected to 10% sodium dodecyl sulfate (SDS)-polyacrylamide gel electrophoresis (PAGE).

### Western Blotting

As described previously ^64, 73^, for quantifying expressions of DMT1, TFR, NCX1-3, Cav-3 or Dab2, and for detection the phosphorylation of cPLA2 and P65, the samples containing 100 μg of protein were added to slab gels. After transferring to PVDF membranes, the samples were blocked by 10% skimmed milk powder for 1 h, and membranes were incubated overnight with the primary antibodies, specific to either DMT1 at a 1:300 dilution, TFR at a 1:200 dilution, NCX1 at a 1:100 dilution, NCX2 at a 1:200 dilution, NCX3 at a 1:150 dilution, Cav-3 at a 1:200 dilution, Dab2 at a 1:100 dilution or β-actin at a 1:1000 dilution. After washing, specific binding was detected by horseradish peroxidase-conjugated secondary antibodies. Images were analysed with an Electrophoresis Gel Imaging Analysis System (MF-ChemiBIS 3.2, DNR Bio-Imaging Systems, Israel). Band density was measured with Window AlphaEase™ FC 32-bit software.

### Monitoring of [Ca^2+^]_i_

For [Ca^2+^]_i_ monitoring and imaging in cultured astrocytes, experiments were run as previously described ^74, 75^. After the pre-treatment with or without inhibitors or siRNA duplex chains, the primarily cultured astrocytes were loaded with 5 μM fluo-4-AM, molecular probes (Thermo Fisher Scientific (Waltham, MA USA)), for 30 min. Fluo-4 signals were visualised by fluorescent microscopy (Olympus IX71, Japan). The fluorescence intensity was normalised to the baseline intensity before stimulation.

### Two-photon *in vivo* Ca^2+^ imaging

As described previously ^75, 76^, adult FVB/N-Tg(GFAP-eGFP)14Mes/J transgenic mice (10 to 12 weeks old) were anesthetized with ketamine (80 mg/kg, i.p.) and xylazine (10 mg/kg, i.p.). Body temperature was monitored using a rectal probe, and the mice were maintained at 37°C by a heating blanket. A custom made metal plate was glued to the skull with dental acrylic cement and a cranial window was prepared over the right hemisphere at 2.5 mm lateral and 2 mm posterior to bregma. The cortical cells were loaded with Ca^2+^ indicator Rhod-2 AM (50 μM, 1 hour). The transcranial window was superfused with artificial CSF. After a stable baseline recording was obtained, Fe^2+^ (100 μM) or Fe^3+^-TF (100 μM) was added for 1 min. Bandpass filters (Chroma) were 540nm/40nm for eGFP and 850nm/70nm for rhod-2 signals. Time-lapse images of astrocytic Ca^2+^ signalling were recorded every five second using FluoView with a custom-built two-photon laser-scanning setup (Nikon AR1, Japan).

### Intracellular Na^+^ measurements

For monitoring intracellular ionized Na^+^ ([Na^+^]_i_) in cultured astrocytes, the measurements were performed as described in ^47^. Primary cultured astrocytes were loaded with 10 μM of Na^+^-sensitive indicator SBFI-AM for 30 min in serum-free medium, with subsequent 1 hour of washout. SBFI was alternatively excited at 340 nm and 380 nm, and the emission was monitored at 500 nm. The SBFI signals were measured by fluorescent microscopy (Olympus IX71, Japan) and expressed as a ratio (R=*F_340_/F_380_*).

### Immunofluorescence

The brain tissue was fixed by immersion in 4% paraformaldehyde and cut into 100 μm slices (see also ^64, 73^). The cultured cells were fixed with 100% methanol at −20 °C. Brain slices or cells were permeabilised by incubation for 1 hour with donkey serum. Primary antibodies against DMT1 or TFR were used at a 1:100 dilution, against glial fibrillary acidic protein (GFAP) was used at 1:200 dilution, and nuclei were stained with markers 4’, 6’-diamidino-2-phenylindole (DAPI) at 1:1000 dilution. The incubation with the primary antibodies were overnight at 4 °C and then donkey anti-mouse or anti-rabbit Cy-2/3 conjugated secondary antibody for 2 h at room temperature. Images were captured using a confocal scanning microscope (DMi8, Leica, Wetzlar, Germany).

### ELISA Assays

Astrocytes were incubated at 37 °C in fresh serum-free culture medium; after the treatment with Fe^2+^/Fe^3+^ or inhibitors, the astrocytes were collected and centrifuged at 10,000×g for 10 min to remove floating cells and/or cell debris at 4°C. To assay the NKA activity, a commercial ELISA kit (abx255202; Abbexa, Cambridge, UK) was used and operated as the protocols, the sensitivity is 0.19 ng/mL, the optical density (OD) was measured at 450 nm and the OD value was normalized by control group. To assay the InsP_3_ concentration ^77^, the supernatant was collected and the concentration of InsP_3_ assayed using a commercial ELISA kit (E-EL-0059c; Elabscience Biotechnology, Wuhan, China), the sensitivity is 10 pg/mL. PGE2 concentrations were measured using a specific ELISA kit (SEKM-0173, Solarbio Life Sciences, Shanghai, China), the assay was performed as the manufacturer’s protocols ^32, 33^.

PGE2 production was evaluated from a standard curve of PGE2, and the released level was finally calibrated by the protein content. The sensitivity of the assay allowed detection is 3 pg/ml. When necessary, the samples were diluted in the assay buffer. The level of AA was determined by using a commercial ELISA kit (E-EL-0051c; Elabscience Biotechnology, Wuhan, China), the sensitivity is 1 ng/mL. The results were normalized by control group and presented as the percentage ^32, 33^.

Astrocytes were incubated at 37 °C in fresh serum-free culture medium; after the treatment with Fe^2+^/Fe^3+^ or inhibitors, the astrocytes were collected and centrifuged at 10,000×g for 10 min to remove floating cells and/or cell debris at 4°C. To determine the NKA activity, a commercial ELISA kit (abx255202; Abbexa, Cambridge, UK) was used and operated as prescribed; the sensitivity is 0.19 ng/mL, the optical density (OD) was measured at 450 nm and the OD value was normalized to the control group. To assay the InsP_3_ concentration ^77^, the cultured cells were placed into lysate and centrifuged at 10,000×g at 4°C for 10 min, the supernatant was collected and the concentration of IP3 assayed using a commercial ELISA kit (E-EL-0059c; Elabscience Biotechnology, Wuhan, China). Preparation of the reference standard and sample diluent, adding working solution, substrate reagent and stop solution in sequence. Determine the OD value with a micro-plate reader at 450 nm, the sensitivity is 10 pg/mL. PGE2 concentrations were measured using a specific ELISA kit (SEKM-0173, Solarbio Life Sciences, Shanghai, China), the assay was performed as the manufacturer’s protocols ^32, 33^. PGE2 production was evaluated, the amounts were calculated from a standard curve of PGE2, and the released level was finally calibrated by the protein content. The sensitivity of the assay allowed detection is 3 pg/ml. When necessary, the samples were diluted in the assay buffer. And the level of AA was assayed by using a commercial ELISA kit (E-EL-0051c; Elabscience Biotechnology, Wuhan, China), the sensitivity is 1 ng/mL. The results were normalized by control group and presented as the percentage ^32, 33^.

### Sorting neural cells through fluorescence activated cell sorter (FACS) and Quantitative PCR (qPCR)

To measure the mRNA for TFR and DMT1, astrocytes expressing fluorescent marker GFP (GFAP-GFP mice) and neurones expressing fluorescent marker YFP (Thy1-YFP mice) were used; we also extracted the cerebral hemispheres tissues from wild type mice. As previously described ^76, 78^, the cells from transgenic mice were used for specific sorting of astrocytes or neurones with FACS. The RNA of the sorted cells and cerebral tissue was extracted by Trizol. Total RNA was reverse transcribed and PCR amplification was performed in a Robo-cycler thermocycler, as per previous description ^73, 76^. Relative quantity of transcripts was assessed using five folds serial dilutions of RT product (200 ng). RNA quantity was normalised to glyceraldehyde 3-phosphate dehydrogenase (GAPDH) and values are expressed as the ratio TFR/GAPDH or DMT1/GAPDH.

### Statistical Analysis

For statistical analysis we used one-way analysis of variance (ANOVA) followed by a Tukey post hoc multiple comparison test for unequal replications using GraphPad Prism 5 software (GraphPad Software Inc., La Jolla, CA) and SPSS 24 software (International Business Machines Corp., NY, USA). All statistical data in the text are presented as the mean ± SD, the value of significance was set at p < 0.05.

## Acknowledgments

This study was supported by Grant No. 81871852 to BL from the National Natural Science Foundation of China, Grant No. XLYC1807137 to BL from LiaoNing Revitalization Talents Program, and Grant No. 20151098 to BL from the Scientific Research Foundation for Returned Scholars of Education Ministry of China. Grant No. 20170541030 to MX from the Natural Science Foundation of Liaoning Province.

## Author Contributions

A.V., M.X. and B.L. designed and supervised the study; M.X., W.G., S.L., G.W., N.X., BN.C., BJ.C., M.J. and W.G. collected the data *in vitro* and analysed the relevant data; SS.L., Z.L., C.D., D.Z. and X.L. performed the experiments *in vivo* and analysed the data; B.L. and A.V. wrote the manuscript.

